# ICE: An imputation framework for detecting cellular senescence using weak single-cell signatures

**DOI:** 10.1101/2025.05.25.656048

**Authors:** Peng Xu, Hantao Zhang, Siyao Zhu, Yimeng Kong

## Abstract

**Background:** Cellular senescence represents a stable cell cycle arrest state that plays critical roles in tissue aging and age-related pathologies. Although single-cell RNA sequencing (scRNA-seq) enables comprehensive profiling of millions of cells in aged tissues, reliably identifying senescent cells remains challenging due to the weak and non-specific expression of their marker genes. Existing computational methods can detect active cell states through gene set scoring, but their performance for senescence genes has not been rigorously evaluated.

**Results:** In this study, we show that senescence genes display weak, non-specific expression patterns across different tissues and are susceptible to dropout events in scRNA-seq. Current scoring approaches suffer intrinsic limitations in single-cell detection using weakly expressed markers. Simply expanding the gene set of weak markers failed to improve detection accuracy. To overcome these limitations, we developed ICE (Imputation-based <u>C</u>ell <u>E</u>nrichment), a computational framework that combines expression imputation with iterative marker refinement. ICE substantially improved detection precision in pancreatic α cells with weakly expressed markers, increasing it from 56% to 98%. Using consensus markers, ICE identified senescent fibro-adipogenic progenitors (FAPs) in muscle tissue, characterized by elevated expression of senescence-associated secretory phenotype (SASP) genes. When applied to individual senescence markers (*CDKN1A/p21, CDKN2A/p16, ATF3*, and *MX1*), ICE successfully identified marker-specific cell populations. These included stressed β cells in aging and type-I interferon-responsive microglia in Alzheimer’s disease (AD).

**Conclusions:** Our study introduces ICE, a rigorous framework for detecting cell states defined by weakly expressed markers at the single-cell level. This tool enables the reliable identification of senescence-associated populations, facilitating a deeper characterization of their heterogeneity and temporal dynamics across diverse human tissues and disease contexts.

## Introduction

Aging is a complex, multifactorial process characterized by the progressive decline of physiological functions across organ systems[1]. At the cellular level, cellular senescence—a state of irreversible cell cycle arrest— plays a central role in organismal aging[2]. This process is triggered by diverse stressors, including DNA damage, telomere attrition, oncogenic activation, and oxidative stress[3]. Key regulators of senescence include cell cycle inhibitors, DNA damage response mechanisms, and metabolic reprogramming. A hallmark of senescent cells is the senescence-associated secretory phenotype (SASP), which involves the secretion of inflammatory cytokines, chemokines, and proteases that disrupt tissue microenvironments and contribute to age-related dysfunction[4, 5].

Despite its significance, studying senescence in human aging remains challenging due to the absence of universal biomarkers[6, 7]. Conventional markers such as β-galactosidase activity, *CDKN1A*/*p21* and *CDKN2A*/*p16* are context-dependent and often overlap with non-senescent processes[8]. For example, *p21* participates in both senescence and oligodendrocyte development, limiting its specificity as a definitive senescence marker[9]. Recent advances have identified novel senescence-associated pathways and gene markers. For instance, multiple studies reported the derepression of LINE-1 retrotransposons and senescence-associated endogenous retroviruses (SA-ERVs), which activate the IFN-I signaling pathway[10-12]. The transcription factor ATF3 drives SA-ERV expression, producing double-stranded RNAs that amplify IFN-I signaling and stabilize the senescent state[13]. ATF3 also remodels chromatin accessibility to promote senescence in endothelial cells and aging tissues, underscoring its pleiotropic role in aging[14, 15]. In the aged mouse brain, microglial IFN-I signaling exacerbates neuroinflammation and neuronal decline, while its ablation mitigates these effects[16]. These findings highlight other non-universal biomarkers, such as *ATF3* and IFN-I signaling, as promising new markers of senescence in human tissues and diseases.

scRNA-seq has transformed aging research by enabling high-resolution analysis of cellular heterogeneity. Studies in model organisms and humans have uncovered senescence-associated transcriptional changes in adipose, muscle, and brain tissues[17-19]. However, detecting senescent cells remains difficult due to the weak and non-specific expression of canonical markers (e.g., *p16, p21*, SASP factors) and inherent technical noise in scRNA-seq. Gene Set Enrichment Analysis (GSEA), originally developed for bulk RNA-seq data, has been adapted to detect aging-associated cells[20, 21]. More recently, methods such as AUCell and UCell have been developed specifically for scRNA-seq to detect active cells using predefined gene markers[22, 23]. However, systematic benchmarking of these methods for weakly expressed senescence markers is lacking.

In this study, we first explored the expression of senescence genes across different tissues and cell types, illustrating their weak and non-specific expression patterns. We then show that current gene set scoring methods, including those designed for bulk and for single-cell data, are intrinsically limited in analyzing weakly expressed markers. To address this limitation, we introduced ICE, an imputation-based analytical framework that enhances detection performance specifically for weak markers. Applying ICE to diverse senescence markers, we identified distinct cell populations in pancreatic β cells and Alzheimer’s disease (AD) microglia. Our method provides a robust platform for investigating senescence heterogeneity and its systemic impacts in aging and age-related diseases at single-cell level.

## Results

### Senescence genes exhibit weak expression in human tissues and cell types

To characterize the expression patterns of senescence-associated genes, we systematically examined senescence marker genes across multiple human tissues and cell types. Using the CellAge database (version 3)[24], we identified 274 senescence-induced genes, including 126 oncogene-induced senescence genes, 65 replicative senescence genes, and 83 stress-induced senescence genes. To assess whether these markers exhibit tissue specificity, we compared their expression levels with tissue-specific genes (tissue-markers) in bulk RNA-seq data from 38 GTEx tissues. To illustrate senescence gene expression, we focused on samples from donors over 50 years of age. The heatmap shows that, unlike tissue-specific markers, senescence markers were expressed non-specifically across diverse tissues (**Fig. 1a**). The median expression of senescence genes ranged between 8.39–8.98 log2(CPM), significantly lower than the median expression of tissue-markers (10.93 log2(CPM); Wilcoxon test, p < 0.01) (**Fig. 1b**). These results suggest that senescence markers are broadly expressed at low levels across tissues, lacking strong tissue specificity.

**Fig. 1:**
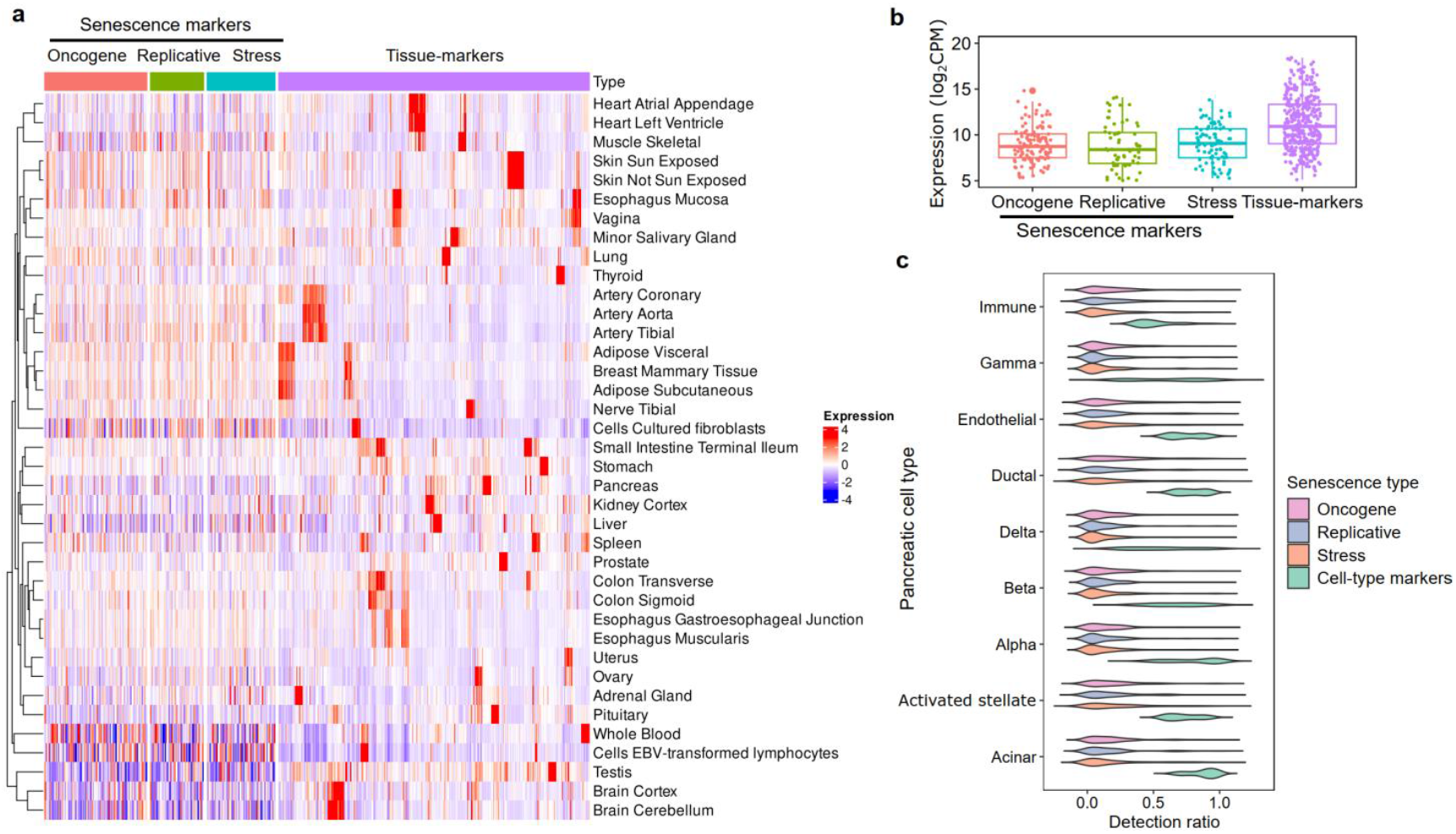
Expression patterns of senescence markers in human tissues and cell types. **a**, Heatmap showing the expression of senescence markers across 38 tissues from the GTEx (v8) database, using data from donors over 50 years old. Rows represent tissues, and columns represent genes grouped by functional categories from the CellAge database. Gene expression values were first averaged across all samples within each tissue and then standardized (z-score) across all tissues. The color scale indicates relative expression, from low (blue) to high (red). **b**, Boxplots of expression distributions for senescence markers (grouped by type) and tissue-specific markers. Boxes denote the interquartile ranges (IQR) (25th–75th percentiles), center lines indicate medians, and whiskers extend to 1.5×IQR. Individual genes are shown as points. **c**, Violin plots depicting detection rates in pancreatic scRNA-seq data. Senescence markers are colored by type.

We next performed single-cell resolution analysis of senescence gene expression using pancreatic scRNA-seq data[25]. To account for technical limitations of scRNA-seq (e.g., dropout events), we calculated detection ratios (1 - dropout rate) for senescence markers. All three senescence marker categories showed low median detection ratios (oncogene-induced: 12.5%; replicative: 10.2%; stress-induced: 8.3%) compared to canonical cell-type markers (71.2%) (**Fig. 1c**). This pattern was consistent across all pancreatic cell types, indicating that senescence markers are weakly expressed and distinct from conventional cell-type markers.

### Benchmarking gene set scoring methods with weak markers

Given the weak expression patterns of senescence genes, we systematically evaluated gene set scoring methods for senescence detection using weak markers. We selected six widely used methods for active cell detection with predefined gene sets, including four bulk-based scoring methods (GSEA[21], GSVA[26], zscore[27], and PLAGE[28]) and two single-cell-based methods (UCell[23] and AUCell[22]). For benchmarking, we selected scRNA-seq datasets from 3 endocrine cell types in the pancreas and 6 major cell types in the brain, all well-characterized and associated with aging-related disorders[25, 29]. Since these datasets lack ground-truth senescent cells, we classified cell-type markers as strong (top 50) or weak (ranks 51–100) based on expression specificity and used the weak markers for evaluation. To further assess robustness, we randomly selected 10–50 marker genes to simulate varying gene set sizes. This benchmarking strategy allowed us to comprehensively evaluate method performance across different levels of marker weakness and gene set sizes.

In addition to the six methods above, we present ICE (Imputation-based <u>C</u>ell <u>E</u>nrichment), a novel approach designed to handle weak markers and dropout effects in scRNA-seq data (**Fig. 2a**). To address technical noise and sparsity, ICE uses MAGIC, a diffusion-based model to smooth cell profiles and impute zero-inflated gene expression[30]. To calculate the enrichment score (ES), ICE embeds GSEA, which performs a weighted random walk along a ranked gene list to measure the overrepresentation of a gene set at the extremes[21]. Unlike conventional methods, ICE provides a binary classification of enriched cells by detecting peaks in the derivative of the ES trend, rather than simply providing numerical scores. To improve robustness for weak markers, ICE iteratively refines gene sets or selects top-correlated genes. This combined imputation and refinement framework ensures reliable enrichment analysis, particularly for low-signal gene sets.

**Fig. 2:**
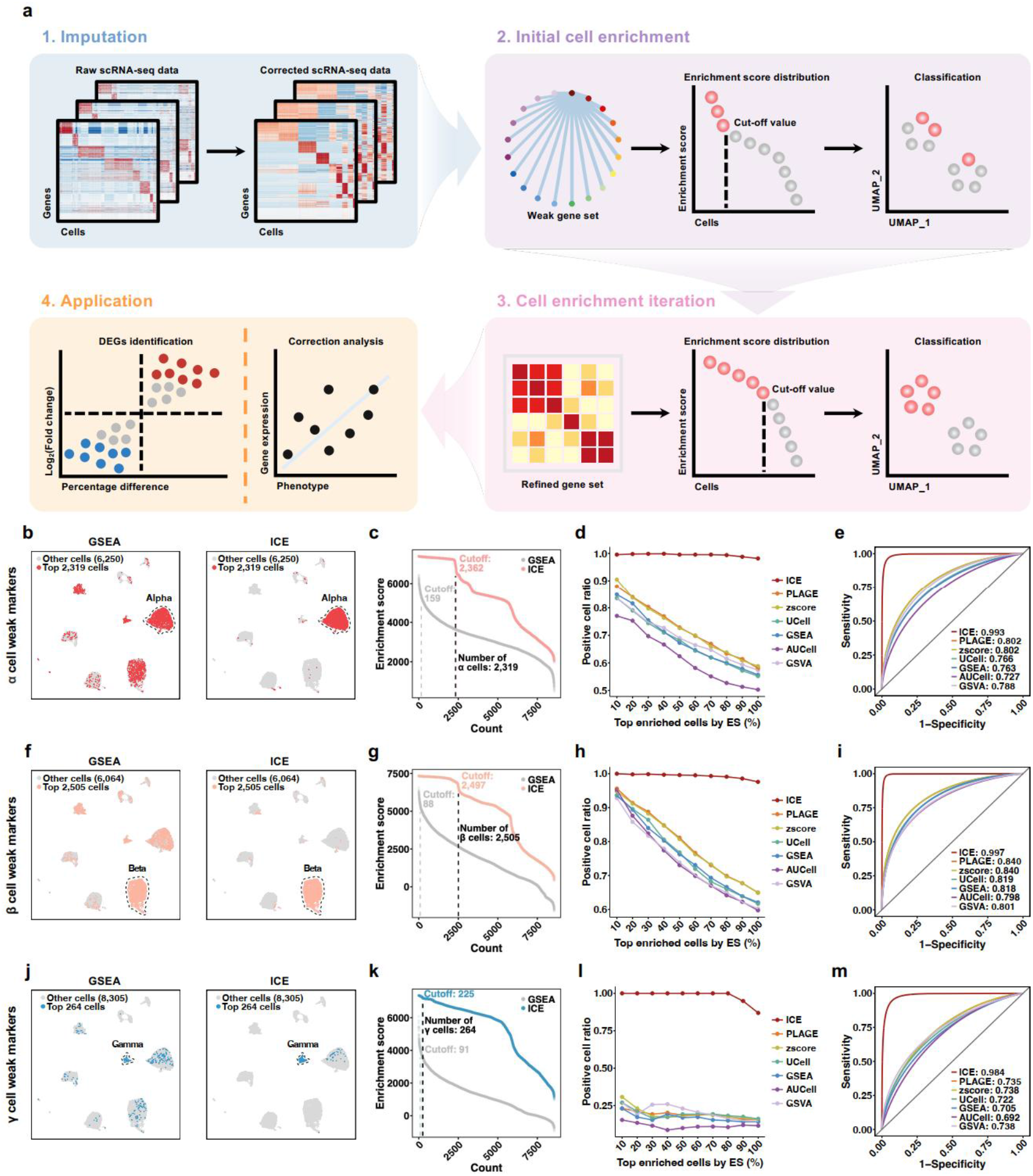
Benchmark of gene set scoring methods using weakly expressed gene markers. **a**, Schematic of the ICE pipeline: (1) denoising and imputation of zero-inflated single-cell data, (2) enrichment score (ES) calculation, (3) significance threshold determination through variation trend analysis, and (4) iterative marker gene refinement to update the ES matrix. **b-e**, Pancreatic α cell detection using 10 weak markers: **b**, UMAP visualization comparing bulk-based GSEA (left) and ICE (right) detection; top 2,319 ES cells (red) versus Louvain-clustered α cells (dashed outline). **c**, ES distribution (ICE: colored; bulk-GSEA: gray) with peak-detection threshold (dotted line). Peaks were detected by derivative analysis of the ES trend. **d**, Precision analysis across ES-ranked cell intervals (10% bins). **e**, ROC curves with AUC values for all methods. **f-i**, Pancreatic β cell evaluation (format consistent with above panels). **j-m**, Pancreatic γ cell assessment (format consistent with above panels).

### ICE outperforms other methods in detecting pancreatic α cells with weak gene markers

Among 8,569 pancreatic cells, we first benchmarked gene set scoring methods for detecting α cells with 10 weak marker genes. Using Louvain clustering as our reference standard, we established a ground truth of 2,319 α cells in the dataset (**Methods**). To evaluate the advantage of ICE’s imputation strategy, we first compared ICE directly with conventional bulk-based GSEA for α cell detection with 10 weak α markers. When analyzing the top 2,319 cells ranked by ES, the bulk-based GSEA approach correctly identified only 1,290 α cells (55.6% precision) (**Fig. 2b**). In contrast, ICE correctly classified 2,276 cells (98.1% precision) and reduced the error rate from 44.4% to 1.9%, demonstrating superior performance.

Further examination of enrichment score distributions revealed critical differences between ICE and the conventional GSEA method. ICE’s peak detection algorithm precisely identified the biological cutoff. The ES values declined abruptly after the top 2,362 cells, closely matching the reference α cell count (**Fig. 2c**). In contrast, traditional GSEA failed to produce a clear separation between α cells and other cell populations in the ES distribution. These results suggest that ICE framework effectively recovers true expression signals that are obscured by dropout events, enabling more accurate enrichment score calculations.

Next, we systematically evaluated seven gene set scoring methods in α cells. To assess marker expression across different abundance levels, we analyzed the ES-ranked cells in cumulative 10% intervals (from top 10% to 100%). ICE demonstrated exceptional robustness, maintaining >98% α-cell identification precision across all expression thresholds (**Fig. 2d**). In contrast, the remaining six methods exhibited expression-level dependence: while achieving reasonable performance (average precision: 84.6%) for the top 10% of cells, their precision markedly declined to 55.9% when analyzing the full cell population. Receiver operating characteristic (ROC) analysis confirmed these findings, showing that ICE achieved near-perfect classification with Area Under the ROC Curve (AUC) value > 0.99 compared to other methods (AUC = 0.73-0.80; **Fig. 2e**). The results demonstrate ICE’s ability to reliably detect cells regardless of marker expression strength.

### ICE demonstrates superior performance in pancreatic β and γ cells

Beyond α cells, ICE demonstrated superior performance in identifying pancreatic β cells with 10 weakly expressed markers. Among 2,505 β cells, traditional GSEA detected only 1,554 (62.0%) with 10 weak markers, whereas ICE correctly identified 2,442 (97.5%) (**Fig. 2f**). The enrichment score (ES) distribution revealed that ICE’s peak detection effectively distinguished β cells from others, while traditional GSEA failed (**Fig. 2g**). ICE maintained high accuracy across varying marker expression levels, whereas other methods declined to 60–70% as ES decreased (**Fig. 2h**). Consistently, ICE achieved an AUC value of 0.99 for β cell detection, outperforming other methods (0.80–0.84) (**Fig. 2i**).

Detecting γ cells, which were rare in the pancreatic dataset (3.0%, n=264), was even more challenging with 10 weak markers. Traditional GSEA identified only 37 (14.0%) γ cells, while ICE detected 227 (86.0%) (**Fig. 2j**). Despite the small population, ICE’s ES distribution allowed peak-based cutoff determination (**Fig. 2k**). ICE exhibited 100% precision in the top 80% of ES-ranked cells, dropping to 86.9% when all cells were included (**Fig. 2l**). In contrast, other methods performed poorly (11.5–16.2%) across all thresholds. ICE’s AUC value (0.98) far surpassed competing methods (0.69–0.74) (**Fig. 2m**).

### Evaluation of ICE’s imputation and marker iteration steps

Comprehensive evaluation of each tool using additional metrics, including recall, F1-score, and Precision-Recall AUC (PR-AUC), revealed that ICE consistently outperformed all other methods across these metrics in α, β, and γ cells (Additional file 1: **Table S1**). DeLong’s test confirmed that ICE’s superior performance in ROC-AUC was statistically significant (*p* < 0.001) when compared to every other method tested (**Table S1**). To test the stability of ICE’s iterative marker refinement step, we partitioned the pancreas dataset into a training set for marker refinement and an independent, held-out test set for performance evaluation. The results showed that when using markers refined on the training data, ICE achieved similarly high performance on the test sets in detecting α, β, and γ cells (Additional file 1: **Fig. S1**).

We next evaluated ICE’s performance with different imputation methods, including MAGIC and SAVER, the top-performing imputation methods from a previous benchmark study[31], along with the widely used deep-learning-based method scVI[32]. Across all tests, ICE consistently outperformed traditional methods without an imputation step, regardless of imputation method used (Additional file 1: **Fig. S2**). While all three methods achieved similar performance for detecting α and β cells, MAGIC achieved better performance in identifying γ cells compared to SAVER and scVI.

To evaluate the individual contributions of the imputation and marker refinement steps, we further performed an ablation analysis using the pancreas dataset with 10 weak markers. Results showed that imputation with MAGIC alone improved the separation of α and γ cells but failed to resolve β cells. However, the subsequent iterative marker refinement step was essential for correctly identifying cutoffs across all three cell types (Additional file 1: **Fig. S3**). Similarly, while imputation with SAVER was insufficient to establish accurate cutoffs for α and β cells, the iterative refinement process enabled correct cell type separation (Additional file 1: **Fig. S3**). These results demonstrate that both imputation and marker refinement steps are critical for ICE’s performance.

### ICE surpasses existing methods in multiple brain cell types

We further validated ICE on 16,929 brain cells, starting with inhibitory neurons (InNs, n=3,724) using 10 weak markers. Traditional GSEA correctly classified only 65.3% (2,430/3,724) of InNs, while ICE achieved 97.8% (3,641/3,724) (**Fig. 3a**). ICE’s ES distribution enabled precise separation of InNs from other cell types (**Fig. 3b**). Regardless of marker strength, ICE maintained >97.7% precision, whereas other methods showed gradual performance decline at low ES (**Fig. 3c**). ICE also attained a near-perfect AUC score (0.99) compared to others (0.88–0.95) (**Fig. 3d**). The advantage of ICE extended to other brain cell types, including astrocytes, endothelial cells, excitatory neurons, oligodendrocytes, and oligodendrocyte precursor cells (**Fig. 3e-n**).

**Fig. 3:**
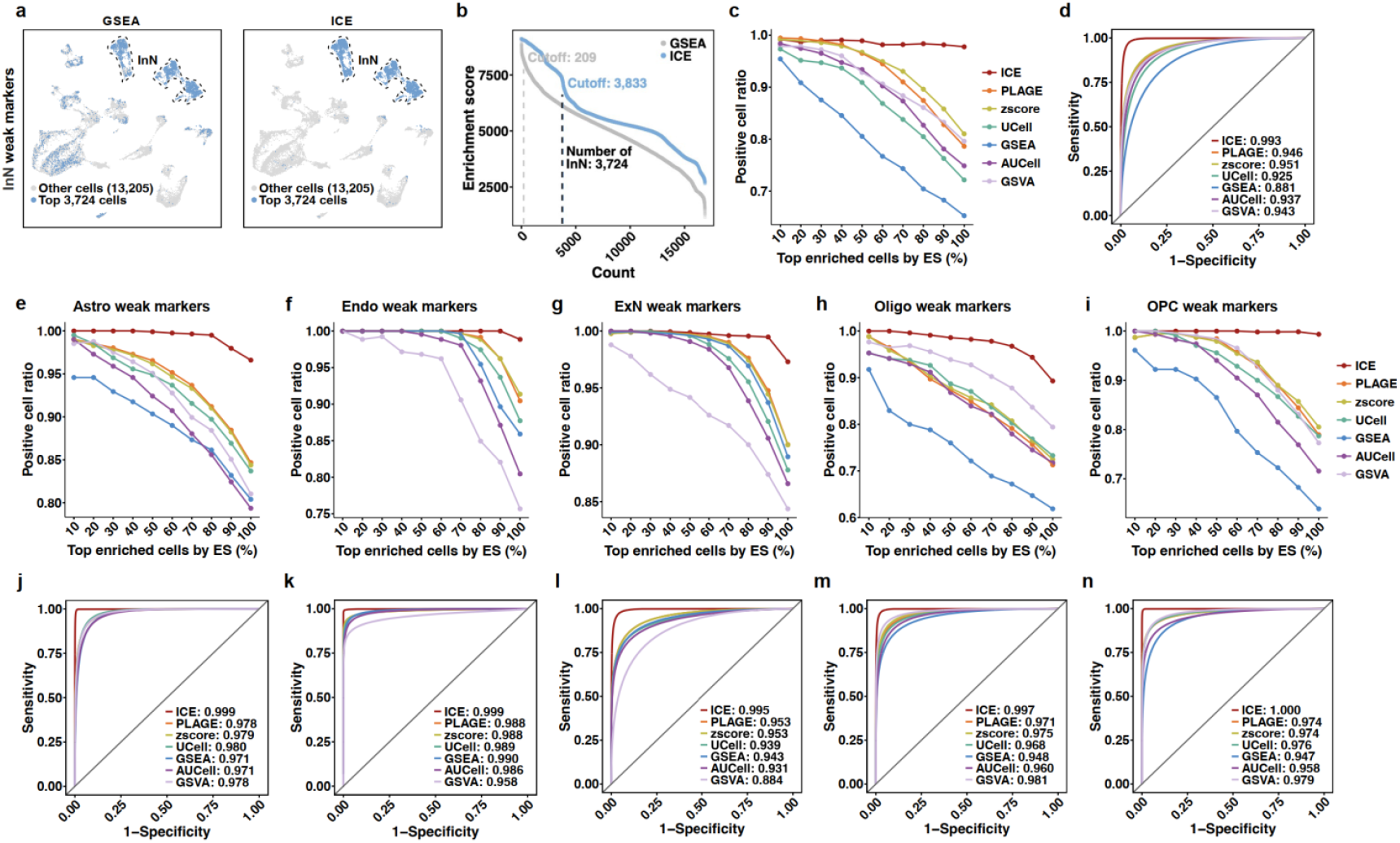
Comparative performance of cell detection methods across brain cell types. **a-d**, Evaluation of inhibitory neuron (InN) detection using 10 weak markers: (**a**) UMAP visualization, (**b**) enrichment score (ES) distribution, (**c**) precision across ES-ranked cells, and (**d**) ROC curves comparing ICE with alternative methods (format consistent with **Fig. 2**). **e-i**, Precision analysis for ES-ranked cells in: (**e**) astrocytes (Astro), (**f**) endothelial cells (Endo), (**g**) excitatory neurons (ExN), (**h**) oligodendrocytes (Oligo), and (**i**) oligodendrocyte precursor cells (OPC). **j-n**, ROC curves with AUC values for each method in corresponding brain cell types (e-i).

### Expanding weak marker sets fails to improve traditional methods

Since traditional methods face inherent constraints in detecting cells using small (10-gene) weak marker sets, we investigated whether increasing the marker panel size could enhance detection performance. We systematically evaluated pancreatic α cell identification across expanding gene sets (20, 30, 40, and 50 markers). ICE maintained consistently superior performance, achieving sustained high precision (∼98.2%) across all marker set sizes **(Fig. 4a-d**). In contrast, other tools showed only modest improvements, with mean precision increasing marginally from 64.5% to 74.9% as marker panels expanded. Importantly, all methods exhibited performance degradation as marker expression strength (ES) decreased. The superior capability of ICE was further corroborated by AUC analysis, where it maintained near-perfect discrimination (∼0.99) regardless of marker set size (**Fig. 4e-h**). Other methods showed substantially lower performance, with the mean AUC value increasing from 0.85 to 0.92, despite the fivefold increase in marker gene number. The results reveal that simply expanding marker set size cannot overcome the fundamental challenge of weak marker strength for traditional methods. However, ICE overcomes this limitation by enhancing the signal from low-expression markers.

**Fig. 4:**
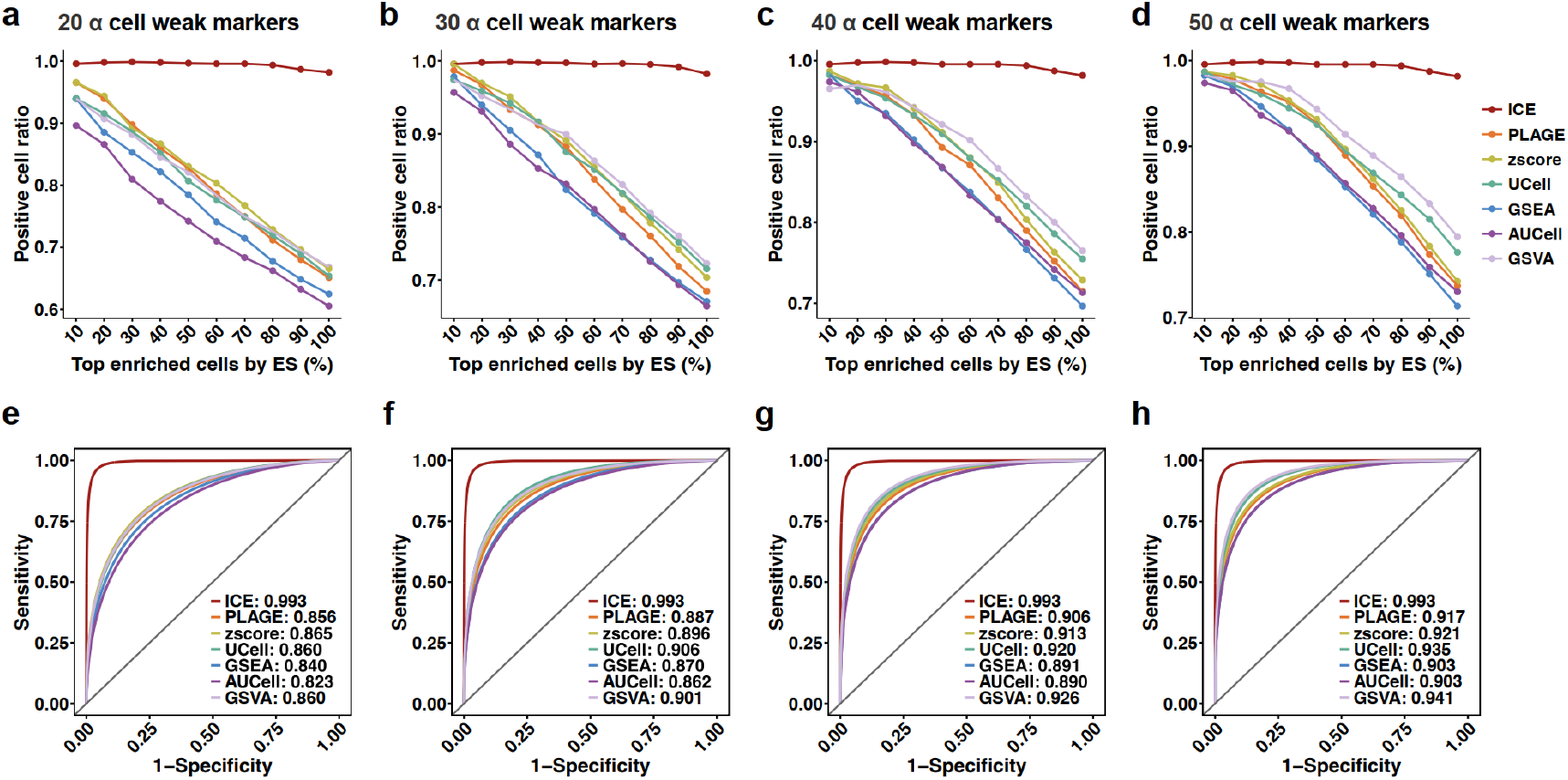
Performance assessment of cell detection methods across varying marker set sizes. **a–d**, Precision of pancreatic α cell detection using weak marker sets of increasing size: (**a**) 20 markers, (**b**) 30 markers, (**c**) 40 markers, and (**d**) 50 markers. **e-h**, Corresponding ROC curves and AUC values for each method: (**e**) 20 markers, (**f**) 30 markers, (**g**) 40 markers, and (**h**) 50 markers.

### ICE effectively identifies senescent cells using consensus markers

Besides the weak cell-type markers, we benchmarked ICE’s performance against established methods in senescent populations using a robust, consensus senescence marker set. Specifically, we used the manually curated SenMayo marker set of 125 senescence genes from the literature[34]. Our evaluation focused on 16,922 fibro-adipogenic progenitors (FAPs) from muscle tissue, a cell type known to contain senescent cells expressing SASP factors and has been used previously for benchmarking[33]. The dataset included 5,729 cells from young (19-27 years old) donors and 11,193 from aged (60-77 years old) donors (**Fig. 5a**). We compared ICE’s performance to GSEA and the specialized tool SenSig[35], which uses the gene signature derived from p16^+^ senescent cells in mouse fibrotic tissue [35].

**Fig. 5:**
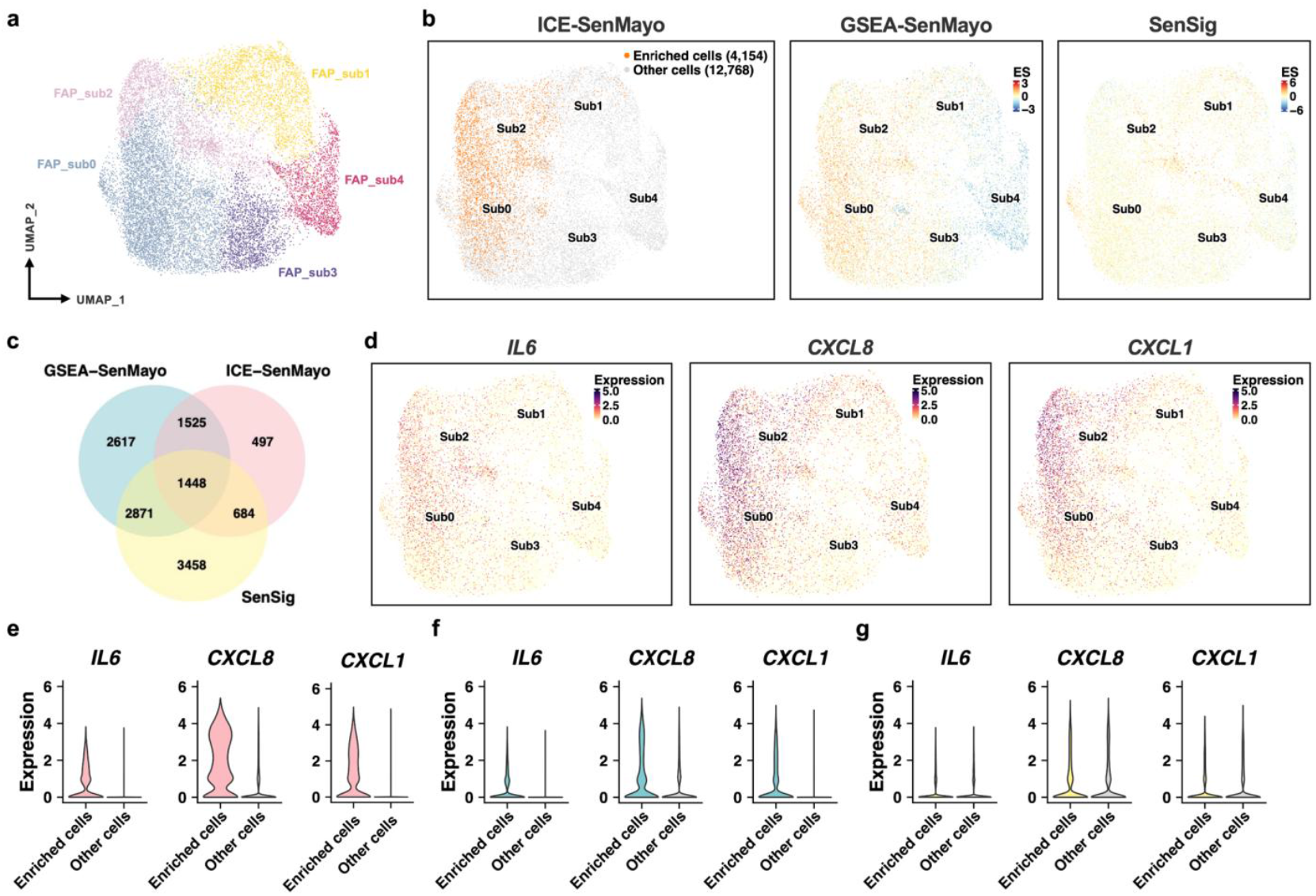
ICE identifies senescent cells in muscle tissue using consensus markers. **a**, UMAP visualization of fibro-adipogenic progenitor (FAP) clusters from muscle tissue. **b**, UMAP plots comparing senescent cell identification across methods. From left to right: cells identified by ICE with the SenMayo marker set (orange), cells colored by GSEA enrichment score (ES) for SenMayo, and cells colored by SenSig ES. **c**, Venn diagram showing the overlap of senescent cells identified by the three methods. The top 50% of ES-ranked cells were regarded as senescent for both GSEA and SenSig. **d**, UMAP plots showing the expression of key senescence-associated secretory phenotype (SASP) genes: *IL6, CXCL8*, and *CXCL1*. **e-g**, Violin plots comparing SASP gene expression in senescent (“Enriched”) versus non-senescent (“Other”) cells. The senescent populations were defined by ICE-SenMayo (**e**), GSEA-SenMayo (**f**), and SenSig (**g**).

Using the SenMayo marker set, ICE initially identified 243 senescent cells, and through iterative refinement, detected a total of 4,154 senescent cells (**Fig. 5b** and Additional file 1: **Fig. S4**). Of these, 99% (4,124/4,154) originated from aged donors. Unlike ICE, which classifies individual cells, GSEA and SenSig provide continuous enrichment scores without a defined cutoff for senescence. Therefore, to enable a direct comparison, we followed a previous benchmarking approach and designated the top 50% of cells ranked by enrichment score as senescent for both methods[33]. A Venn diagram revealed that 3,657 (88%) of the cells identified by ICE were also classified as senescent by GSEA or SenSig (**Fig. 5c**). Meanwhile, GSEA and SenSig each identified additional non-overlapping cells, reflecting tool-specific variability.

Notably, ICE-identified cells exhibited highly expressed SASP genes, including *IL6, CXCL8*, and *CXCL1* (**Fig. 5d**), which are strongly linked to muscle atrophy and fibrosis[33, 36]. Differential gene expression analysis further confirmed that SASP genes were highly expressed in ICE-identified senescent cells compared to non-senescent cells (**Fig. 5e-g**). In contrast, this expression difference was less pronounced for cells identified by GSEA and entirely absent for those identified by SenSig. These findings demonstrate that ICE’s multi-marker approach effectively identifies senescent cells with key functional phenotypes, such as a robust SASP.

### ICE reveals aging-associated *ATF3* activity in β cells

Given ICE’s superior performance, we employed it to detect senescence in human pancreatic β cells (n = 7,528) across different ages[37](**Fig. 6a**). Our evaluation focused on four senescence markers: the canonical markers *CDKN1A* and *CDKN2A*, as well as two novel candidates, *ATF3*, and *MX1* (an interferon-I pathway gene[38]). For each marker, ICE identified the top 20 correlated genes to define marker-specific signatures. Using these signatures, ICE identified 496 (*CDKN1A*), 632 (*CDKN2A*), 154 (*ATF3*), and 379 (*MX1*) marker-associated cells (**Fig. 6b**).

**Fig. 6:**
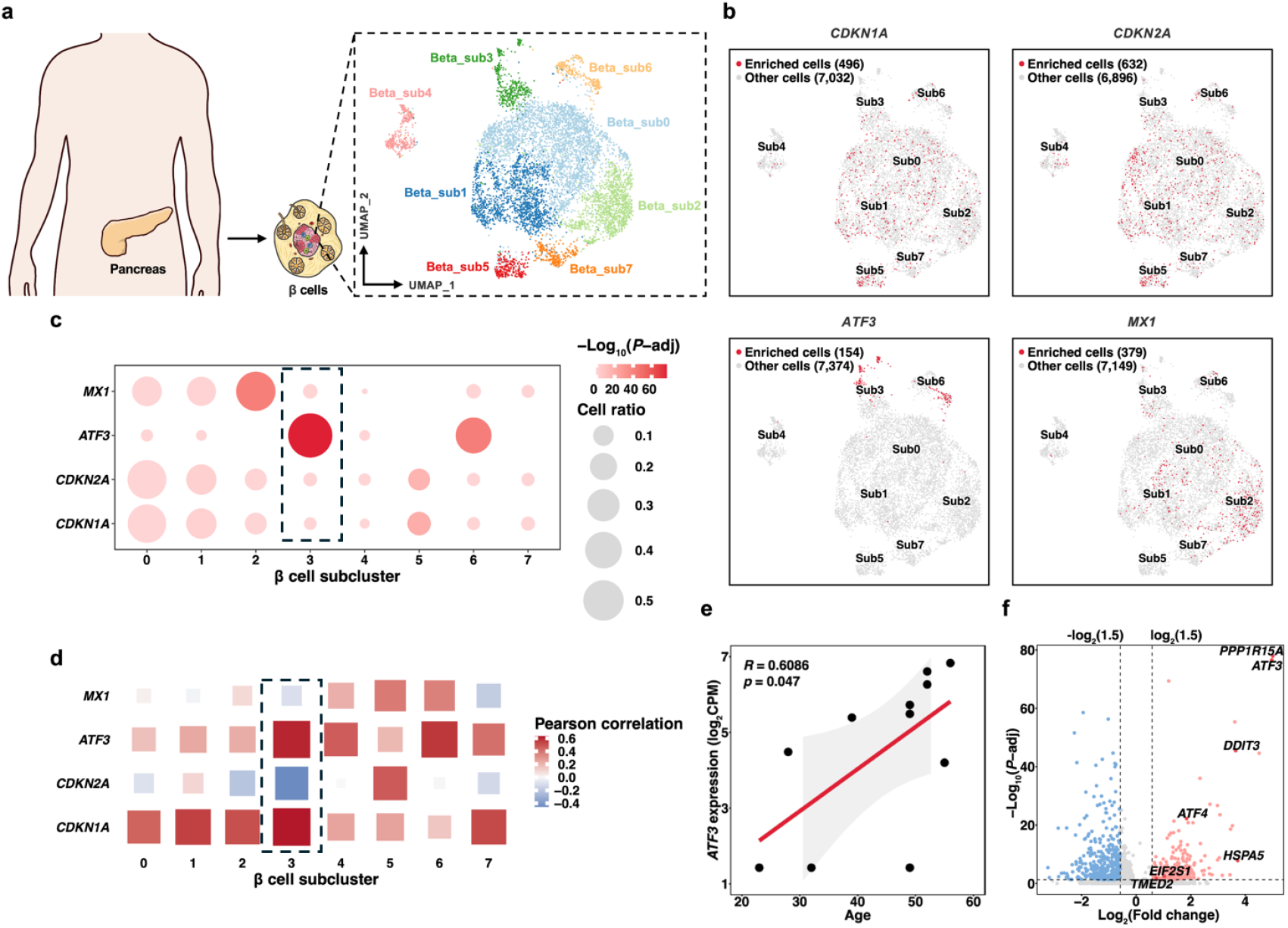
Identification of senescence marker-associated β cell subpopulations using ICE. **a**, UMAP visualization of pancreatic β cell heterogeneity, revealing distinct subclusters. **b**, ICE-based detection of senescence marker-associated cells (red) versus unassociated populations (gray). Gene sets for ICE analysis were constructed by selecting the top 20 correlated genes for each senescence marker. **c**, Cluster-specific enrichment patterns of senescence-associated cells. Circle color: significance level from Fisher’s exact test. Circle size: proportion of enriched cells per subcluster. **d**, Age-related expression dynamics of senescence markers across β cell subclusters. **e**, Pseudobulk analysis in *ATF3*-associated cells confirming positive correlation between *ATF3* expression and donor age. **f**, Volcano plot of differentially expressed genes in *ATF3*-associated β cells. Upregulated genes associated with UPR pathway are labeled. Pink: up-regulated genes; Blue: down-regulated genes.

While *CDKN1A, CDKN2A* and *MX1* displayed diffuse expression patterns, *ATF3*-associated cells were specifically enriched in subclusters 3 and 6 (**Fig. 6b,c**). This suggests that *ATF3*-associated cells formed a distinct subpopulation of β cells. Pseudobulk analysis in subcluster 3 showed that *ATF3* and *CDKN1A* expression increased with age, while *MX1* and *CDKN2A* decreased (**Fig. 6d**). In all *ATF3*-associated cells, expression of *ATF3* showed a positive correlation with the donor age (**Fig. 6e**). Interestingly, differential gene expression analysis revealed that *ATF3*-associated cells exhibited an upregulation of pro-apoptotic unfolded protein response (UPR)-related genes (**Fig. 6f**). Specifically, *DDIT3* (*CHOP*) functions as a key mediator of ER stress-induced apoptosis[39] and *HSPA5* (*GRP78*/*BiP*) encodes a master regulator of the UPR[40]. These findings indicate that *ATF3*-associated β cells become activated during aging and participate in chronic ER stress, UPR activation, and increased apoptosis susceptibility.

### ICE uncovers IFN-I-active microglia linked to AD

IFN-I responsive microglia play a role in neurodegenerative disorders and contribute to neuronal engulfment during development and disease[16, 41]. Therefore, we applied ICE to investigate the link between IFN-I-active microglia and AD[42]. Unsupervised clustering of 76,346 microglia from human AD brains revealed 13 distinct subclusters (**Fig. 7a**). ICE identified 3,212 (*CDKN1A*), 779 (*CDKN2A*), 1,635 (*ATF3*), and 927 (*MX1*) marker-associated cells, each displaying unique expression patterns (**Fig. 7b**). Notably, *MX1*-associated microglia localized specifically to subcluster 5, while *CDKN1A* and *CDKN2A* were restricted to subclusters 2 and 4 (**Fig. 7c**).

**Fig. 7:**
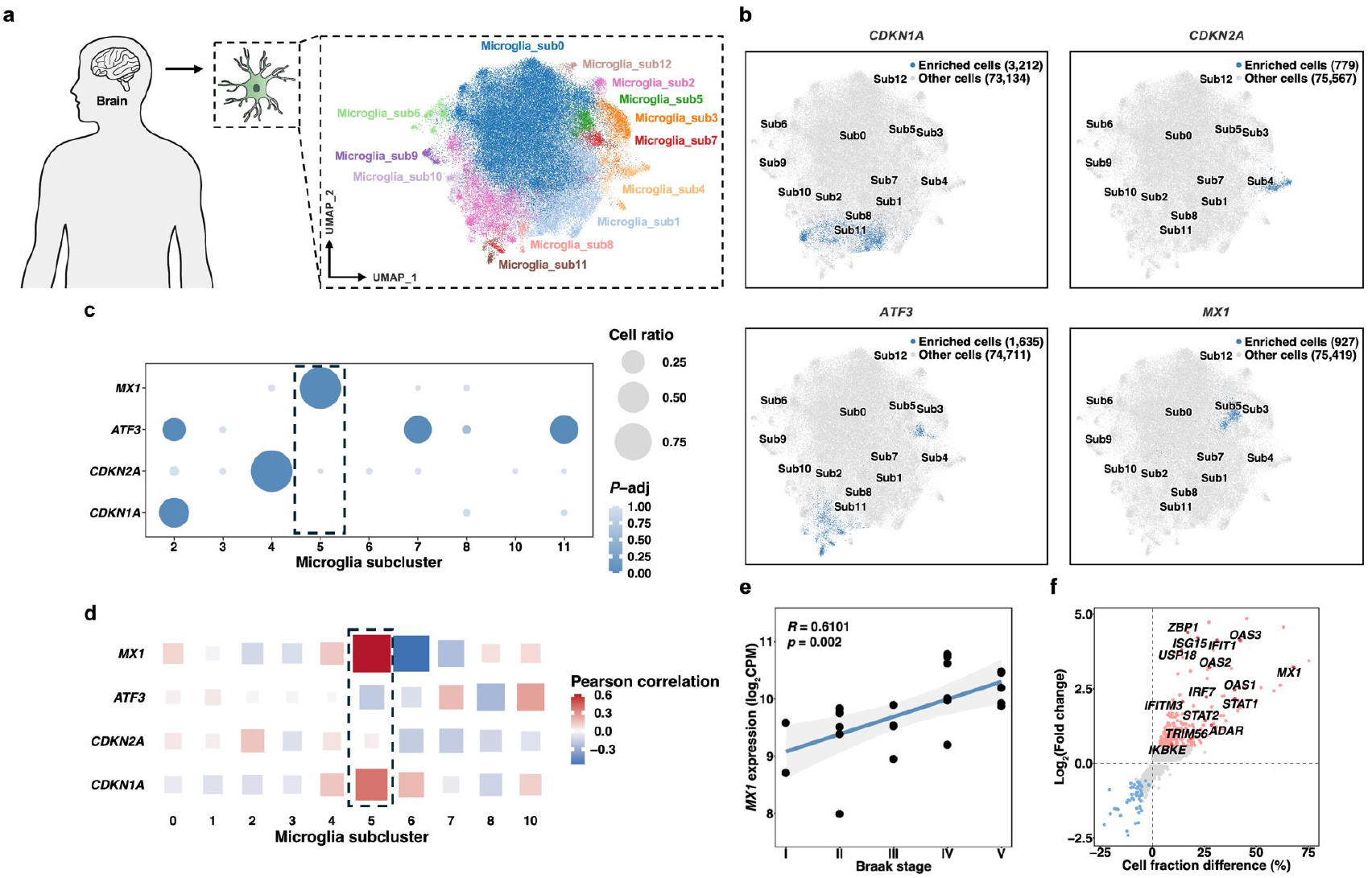
ICE reveals senescence marker-associated microglia subpopulations in AD. **a**, UMAP projection of microglia heterogeneity in AD brains, showing distinct transcriptional subclusters. **b**, Detection of senescence marker-associated microglia (red) versus unassociated populations (gray). Gene sets for ICE analysis were constructed by selecting the top 20 correlated genes for each senescence marker. **c**, Subcluster enrichment patterns of senescence-associated cells. Circle color: significance level from Fisher’s exact test. Circle size: proportion of enriched cells per subcluster. **d**, Expression correlation of senescence markers with AD Braak stage across different subclusters. **e**, Pseudobulk analysis demonstrating positive correlation between *MX1* expression and AD neuropathological progression (Braak stage). **f**, Volcano plot of differentially expressed genes in *MX1*-associated microglia. Upregulated genes associated with IFN-I signaling pathway are labeled. Pink: up-regulated genes; Blue: down-regulated genes.

In subcluster 5, *MX1* and *CDKN1A* expression correlated positively with AD progression (Braak stage), whereas *ATF3* exhibited a mild negative correlation (**Fig. 7d**). Further analysis confirmed a significant positive association between *MX1* expression and Braak stage across all *MX1*-associated cells (**Fig. 7e**). These cells also showed upregulation of key regulators in IFN-I signaling genes, including *STAT1, STAT2, OAS1*, and *OAS2* (**Fig. 7f**). These findings demonstrate ICE’s ability to precisely detect cell populations enriched for senescence markers, highlighting a link between IFN-I-active microglia and AD pathogenesis.

## Discussion

The emergence of single-cell profiling technologies has revolutionized aging research and provided unprecedented insights into senescent cell states. However, the reliable identification of senescent cells remains a significant challenge due to the weak expression of senescence biomarkers. This limitation underscores the urgent need for novel computational approaches to accurately detect senescent cells with subtle molecular signatures.

In this study, we first established a benchmark to evaluate the performance of existing tools in detecting cells based on weakly expressed gene sets. Our analysis revealed critical shortcomings in current gene set scoring methods: as marker expression strength decreased, detection accuracy declined across multiple cell types in the pancreas and brain. For example, in pancreatic γ-cells, precision dropped to as low as 11.5% when markers were particularly faint. Notably, increasing the number of marker genes did not substantially improve performance, suggesting that the primary limitation lies in the inherent weakness of the markers rather than the size of the gene set.

To overcome these challenges, we developed ICE, a computational framework that integrates two key strategies: (1) imputation to recover missing or weak expression signals and (2) iterative refinement of gene sets to enhance marker specificity. Extensive benchmarking demonstrated that ICE significantly outperformed existing methods. In pancreatic α cells, ICE achieved over 98% precision and a near-perfect AUC even with weak markers, whereas conventional methods performed poorly, with precision rates of 50-60%. ICE showed similar superior performance in identifying diverse cell types in the pancreas and brain tissues. These results highlight the importance of expression data imputation and dynamic gene set optimization for reliable cell detection in scRNA-seq data.

Beyond single weak cell-type markers, we also benchmarked ICE against established methods using a robust, consensus senescence marker set. Using the manually curated SenMayo markers[34], ICE identified a senescent FAP population found almost exclusively in aged donors. A key advantage of ICE is its ability to provide a clear classification cutoff. In contrast, methods like GSEA and SenSig only provide numeric enrichment scores, requiring an arbitrary threshold (e.g., top 50% of cells) for classifying senescent cells [33]. The senescent cells identified by ICE showed stronger expression of key SASP genes compared to those identified by GSEA. Moreover, cells with high SenSig scores showed no enrichment for SASP expression. This suggests that the SenSig gene signature, derived from a mouse fibrotic model [35], may not be directly applicable to identify senescent cells in human muscle FAPs.

Applying ICE to key senescence markers (*CDKN1A, CDKN2A, ATF3*, and *MX1*), we uncovered distinct cell populations enriched for these markers and explored their functional implications. In pancreatic β-cells, ICE identified a subpopulation with elevated *ATF3* expression, which exhibited heightened unfolded protein response (UPR) pathway activity, marked by *DDIT3* upregulation. Prior studies have shown that *DDIT3* deletion in β cells mitigates ER stress and prevents age-associated metabolic dysfunction in mice[39]. Thus, the *ATF3*-positive β cells detected by ICE may represent a senescent population under chronic ER stress during aging.

Similarly, in the context of neurodegenerative disease, ICE revealed a subset of interferon-responsive microglia enriched in late-stage Alzheimer’s disease (AD). These cells, characterized by high *MX1* expression, displayed robust IFN-I signaling activity, which has been linked to AD progression. Given that IFN-I signaling can be triggered by damage-associated nucleic acids, these microglia may be activated by amyloid fibrils containing such molecules[43]. Supporting this notion, blocking IFN-I signaling has been shown to improve cognitive and synaptic function in AD models[44]. These findings suggest that *MX1*-expressing microglia represent a chronically inflamed state associated with AD pathology.

The results from the four senescence markers highlight the remarkable heterogeneity in human tissues. We observed minimal overlap among them, with each labeling a distinct cellular population. In aged β-cells, *ATF3* consistently marked a subpopulation associated with age-related activation of the UPR, a phenotype closely linked to stress-induced senescence. In contrast, classical markers such as *CDKN1A* and *CDKN2A* exhibited more variable expression patterns. Moreover, the rare co-expression of *ATF3* and *MX1* in both β-cells and microglia suggests that their transcriptional regulation is highly context-dependent. This evidence indicates that “senescence” is not a uniform state, but rather a spectrum of diverse phenotypes. It should be investigated with careful consideration of marker selection and biological context. Therefore, there is a pressing need to develop more robust, context-specific consensus marker panels for different tissues and disease conditions.

For practical applications, the enrichment scoring and marker refinement steps of ICE are computationally efficient and typically complete within minutes. The primary bottleneck lies in the initial imputation step. While MAGIC performs efficiently on small-to medium-sized datasets, its computational cost increases quadratically (*O*(*N*^2^)). For very large datasets (>100,000 cells), we recommend applying ICE to cell type– specific subsets. In addition, we provide a step-by-step ICE framework that can be coupled with different imputation methods, including more scalable imputation algorithms such as scVI.

In summary, our study introduced ICE, a computational framework that enhances cell-state identification through imputation and iterative marker refinement. By enabling high-resolution dissection of senescence heterogeneity, ICE offers novel insights into the spatiotemporal dynamics of aging across tissues. Moreover, the ICE framework is broadly applicable to detect diseased cells with weakly expressed markers, opening new avenues for studying cellular dysfunction in diverse pathological contexts.

## Methods

### Overview of ICE

ICE is a single-cell gene set enrichment analysis method designed to identify cells significantly enriched for a predefined gene set. The method integrates imputation, iterative refinement, and derivative-based cutoff estimation to enhance the accuracy and robustness of enrichment detection, particularly in the presence of sparse or weak marker genes. ICE requires two main inputs: a gene expression profile of single cells and a predefined set of genes (e.g., a gene set of interest). Below, we outline the key steps of the ICE algorithm:

#### 1. Imputation of Single-Cell Data

The first step in ICE involves rectifying the single-cell gene expression data using the MAGIC algorithm[30], which was ranked as one of the most accurate imputation methods in an extensive benchmark study[31]. The imputation begins by constructing a k-Nearest Neighbor (kNN) graph, where each cell is connected to its k-nearest neighbors based on Euclidean distance to capture cell-to-cell similarities. Next, an affinity matrix is built using a Gaussian kernel to transform these distances into similarity scores, weighted by a parameter to control the spread of the similarity function. Finally, a diffusion process is applied across the kNN graph, propagating expression values based on cell affinities to smooth the data, impute missing values, and reduce noise. The resulting imputed matrix provides a more complete and accurate representation of cellular gene expression profiles for downstream analyses.

#### 2. Calculation of Enrichment Scores (ES)

For each cell, ICE computes an ES that reflects the expression activity of the predefined gene set. In this step, single-sample Gene Set Enrichment Analysis is applied to calculate enrichment scores for predefined gene sets on a per-cell basis [26]. The process begins by ranking genes within each cell based on their expression levels, creating an ordered list from highest to lowest expression. For each gene set, an enrichment score is then computed using a running-sum approach, which assesses the positions of genes within the ranked list to capture their relative contribution to the gene set’s activity. This results in a matrix of enrichment scores, where each entry represents the activity level of a specific gene set in a given cell, providing a comprehensive view of gene set expression across the cell population and enabling insights into cellular states and pathways.

#### 3. Derivative-Based Cutoff Estimation

The ES values are arranged in descending order, and forward differences are computed to approximate the derivatives of the ES trend. The pracma package (v2.4.4) in R was used to calculate the cutoff value from the initial peak in the derivative trend. This cutoff distinguishes cells significantly enriched for the gene set from those that are not.

#### 4. Iterative Refinement via Differential Expression Analysis

Based on the initial cell classification, ICE performs differential expression analysis using the Seurat package[45]. The resulting positive markers are used to refine the discovery of enriched cells. The top n positive marker genes (where n is a user-defined variable) are chosen to replace the predefined marker list. The enrichment scores and cutoff values are then recalculated iteratively to optimize the identification of cells active in the predefined gene set.

The output of ICE not only includes the enrichment scores derived from denoised gene expressions but also provides a clear cutoff for identifying significantly enriched cells.

### Benchmarking different gene set scoring methods

The performance of ICE was systematically benchmarked against four bulk-based scoring methods (GSEA, GSVA, zscore, and PLAGE) and two single-cell-based scoring methods (UCell and AUCell). GSEA evaluates the distribution of predefined gene sets within a ranked list of genes to identify significant biological pathways[21]. GSVA transforms gene expression data into gene set enrichment scores, enabling pathway-level analysis across samples[26]. The z-score method calculates the standardized difference between the mean expression of a gene set and the background, providing a simple yet effective enrichment metric[27]. PLAGE uses singular value decomposition to model gene set activity as latent variables[28]. On the single-cell level, UCell computes a rank-based enrichment score by assessing the distribution of gene set members within the ranked expression profile of each cell[23]. AUCell calculates the Area Under the Curve (AUC) for the recovery curve of gene set members, quantifying their enrichment in individual cells[22].

Three single-cell RNA-seq datasets from human pancreas, muscle, and brain tissues were used for benchmarking analysis. The pancreas and muscle datasets were downloaded from the Gene Expression Omnibus (GEO) under accession numbers GSE84133[25] and GSE268953[33], respectively. The brain dataset was obtained from the PsychENCODE Consortium[29]. All datasets were preprocessed using the Seurat pipeline, including normalization with the SCTransform method, dimensionality reduction via PCA, integration using Harmony, and clustering with the Louvain algorithm at a resolution of 0.2. UMAP was employed for visualization. Cell type annotations were assigned as follows: cell types in the pancreas dataset were predicted using the Azimuth reference mapping tool; the muscle dataset was annotated using established gene markers; and annotations for the brain dataset were refined by assigning the most frequent cell-type label within each cluster. To identify marker genes for each cell type, differential expression analysis was performed using the Wilcoxon rank-sum test.

To evaluate the performance of the methods, three endocrine cell types (α, β, and γ) were analyzed in the pancreas dataset, while six cell types (inhibitory neurons, astrocytes, endothelial cells, excitatory neurons, oligodendrocytes, and oligodendrocyte precursor cells) were chosen from the brain dataset. To assess the impact of marker strength and sparsity, two categories of marker gene sets were defined for each cell type: strong markers comprised the top 50 most differentially expressed genes, and weak markers included the subsequent 50 genes. These differentially expressed genes were identified using the Wilcoxon test during cell-type marker analysis, ranked by their adjusted p-values. Additionally, random subsets of 10, 20, 30, 40, and 50 markers were selected from each category to evaluate the methods under varying marker sparsity conditions. For each cell type, the clustering labels were used as ground truth annotations. Precision was calculated as the proportion of correctly classified cells relative to the total number of identified cells based on the ranked ESs. To further evaluate performance, Receiver Operating Characteristic (ROC) analysis was conducted. The ROC curve was constructed using the “roc” function from the “pROC” package in R, with the true positive rate (TPR) and false positive rate (FPR) calculated based on the normalized ES scores. The overall performance of each method was quantified using the Area Under the Receiver Operating Characteristic Curve (ROC-AUC). To determine whether the differences in ROC-AUC values between methods were statistically significant, we applied DeLong’s test. For a more comprehensive assessment, we also calculated recall, F1-score, and Precision-Recall AUC (PR-AUC) using the “yardstick” package in R.

### RNA-seq analysis of GTEx database

Raw RNA-seq count data were obtained from the GTEx database (v8). Tissues with fewer than 50 samples were excluded, resulting in 14,956 samples from 38 tissues derived from 948 non-diseased donors aged 20– 80 years. For each tissue, genes with expression levels > 1 log2CPM (counts per million) in at least 25% of samples were retained for downstream analysis. Read counts were normalized using the trimmed mean of M-values (TMM) method to correct for library size differences[46]. Normalized expression values were log2-transformed. Tissue-specific differentially expressed genes (DEGs) were identified using the limma package (moderated t-test) [47]. Genes with |log2FC| > 1 and adjusted P-value < 0.05 were classified as significant DEGs and tissue-specific markers.

### Enrichment analysis with single gene markers

To evaluate ICE using single gene markers, we performed correlation analysis based on imputed gene expression data. For each target gene, the top n most correlated genes (including the target gene itself) were selected to construct a marker gene set for ICE. In this study, we set n = 20 as part of our exploratory analysis. Two single-cell RNA-seq datasets related to aging and AD were analyzed: stressed β cells and type-I interferon (IFN-I)-responsive microglia. The β cell dataset (GSE114297) was obtained from GEO and consisted of islet samples from 12 donors without diabetes[37]. After preprocessing using the Seurat pipeline (normalization, clustering, and cell type annotation), the β cell population was selected for ICE analysis. The microglia dataset was obtained from the Synapse database (syn52293417) and included nuclei sequencing data from the prefrontal cortex of human individuals with varying degrees of AD pathology and cognitive impairment[42]. The immune cell count matrix was directly downloaded from the database, and the microglia population was isolated for downstream ICE analysis.

## Statistics and Reproducibility

The performance of different methods was evaluated using precision and ROC analyses. The AUC was computed by the “pROC” package in R to quantify the performance of each method. Enrichment of target cells within cell subclusters was assessed using Fisher’s exact test via the “ClusterProfiler” package. Correlations between gene expression and donor age or AD pathology (Braak stage) were calculated using pseudobulked counts for each donor, generated with the “AggregateExpression” function in Seurat. Pearson correlation was used for these analyses. Differential expression analysis to identify cell-type markers was performed using the Wilcoxon test from the Seurat pipeline.

## Declarations

### Ethics approval and consent to participate

Not applicable.

### Consent for publication

Not applicable.

### Data availability

All datasets used for benchmarking are publicly available. The pancreas and muscle datasets were obtained from the Gene Expression Omnibus (GEO) under accession numbers GSE84133 and GSE268953, respectively. The brain dataset was acquired from the PsychENCODE Consortium. The stressed β cell dataset was retrieved from GEO under accession GSE114297, and the AD microglia dataset was obtained from the Synapse database (syn52293417). The R implementation of the ICE package is available on GitHub (https://github.com/pxulab/ICE). The source codes associated with this study have been deposited in Zenodo (https://doi.org/10.5281/zenodo.17211836).

### Authors’ contributions

PX and YK conceived the concept and designed the study; PX and HZ performed the data analyses, prepared the figures, and wrote the paper; SZ participated in the data analysis. All the authors participated in the discussion and interpretation of the results, reviewed and revised the paper.

## Acknowledgements

This work was primarily supported by the National Natural Science Foundation of China (32570780 to PX and 32470666 to YK), the Interdisciplinary Research Program for Young Scholars (2024FGC1004), the Fundamental Research Funds for the Central Universities (2242025F10004) and start-up fund (RF1028624054 to PX and RF1028623368 to YK) from Southeast University.

## Supplementary information

**Additional file 1:** Supplementary Table S1 and Figures S1–S4 with corresponding legends.

