## Supplementary material for "ICE: An imputation framework for detecting cellular senescence using weak single-cell signatures": Figs. S1-S4 and Table S1

**Table S1. Performance of different scoring methods for cell detection using 10 weak markers**

| Method | Gene set | Recall | F1-score | PR-AUC | DeLong's Test (vs. ICE) |
|---|---|---|---|---|---|
| ICE | α cell weak markers | 0.981 | 0.981 | 0.993 | – |
| | β cell weak markers | 0.975 | 0.975 | 0.992 | – |
| | γ cell weak markers | 0.860 | 0.860 | 0.895 | – |
| PLAGE | α cell weak markers | 0.580 | 0.580 | 0.174 | $p < 0.001$ |
| | β cell weak markers | 0.649 | 0.649 | 0.178 | $p < 0.001$ |
| | γ cell weak markers | 0.155 | 0.155 | 0.019 | $p < 0.001$ |
| zscore | α cell weak markers | 0.587 | 0.587 | 0.630 | $p < 0.001$ |
| | β cell weak markers | 0.649 | 0.649 | 0.721 | $p < 0.001$ |
| | γ cell weak markers | 0.155 | 0.155 | 0.098 | $p < 0.001$ |
| UCell | α cell weak markers | 0.551 | 0.551 | 0.583 | $p < 0.001$ |
| | β cell weak markers | 0.616 | 0.616 | 0.685 | $p < 0.001$ |
| | γ cell weak markers | 0.163 | 0.163 | 0.103 | $p < 0.001$ |
| GSEA | α cell weak markers | 0.556 | 0.556 | 0.589 | $p < 0.001$ |
| | β cell weak markers | 0.620 | 0.620 | 0.681 | $p < 0.001$ |
| | γ cell weak markers | 0.140 | 0.140 | 0.085 | $p < 0.001$ |
| AUCell | α cell weak markers | 0.503 | 0.503 | 0.527 | $p < 0.001$ |
| | β cell weak markers | 0.597 | 0.597 | 0.658 | $p < 0.001$ |
| | γ cell weak markers | 0.117 | 0.117 | 0.070 | $p < 0.001$ |
| GSVA | α cell weak markers | 0.572 | 0.572 | 0.605 | $p < 0.001$ |
| | β cell weak markers | 0.601 | 0.601 | 0.660 | $p < 0.001$ |
| | γ cell weak markers | 0.144 | 0.144 | 0.111 | $p < 0.001$ |

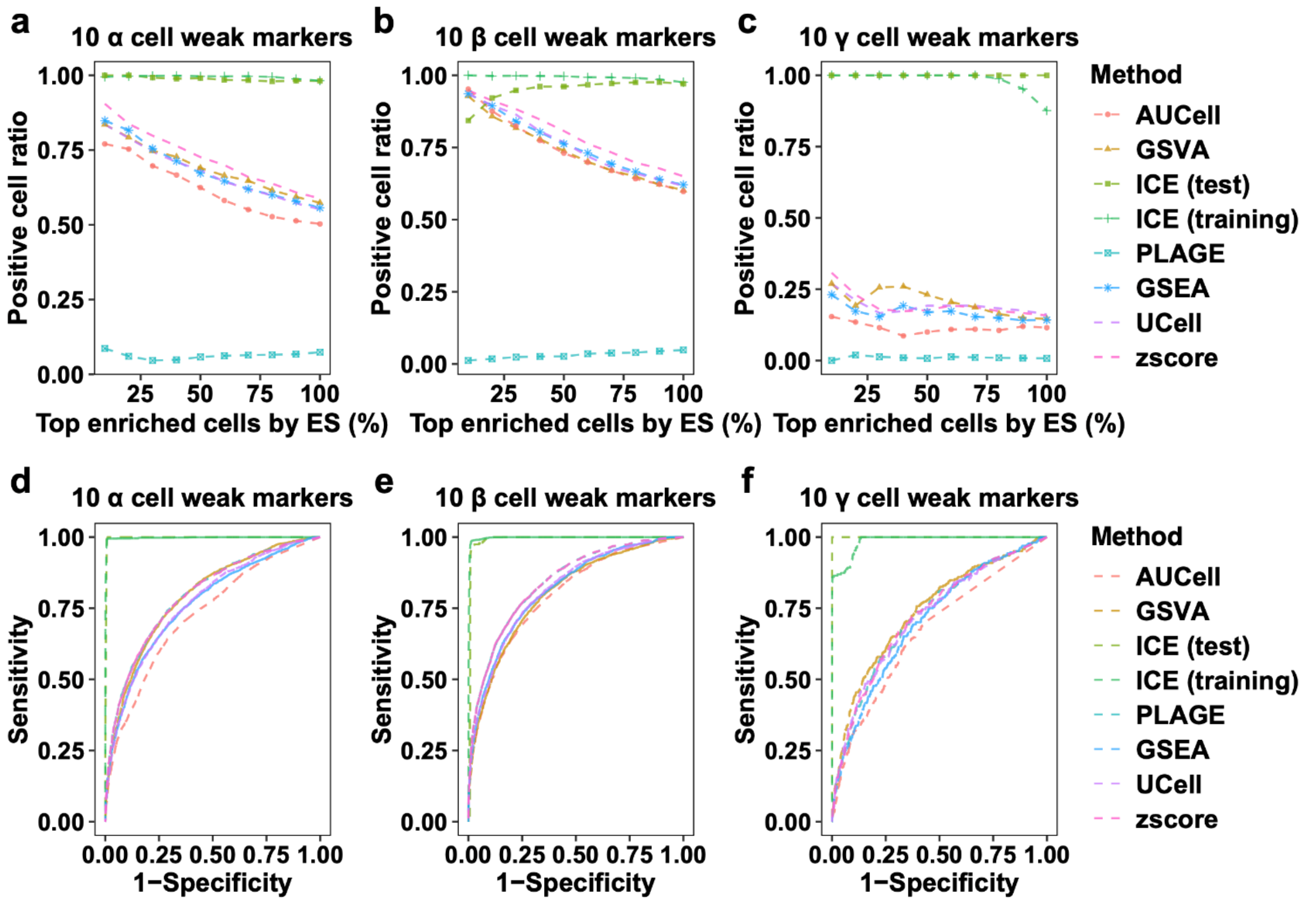


**Fig. S1 | ICE performance is robust and generalizable to independent test data.**

**a-c**, Precision plots across ES-ranked cell intervals for α, β, and γ cells from the pancreas dataset. The analysis compares multiple gene set scoring methods. ICE performance was assessed on the training dataset (80% of cells) and on a held-out independent test dataset (20% of cells) using markers refined from the training data.

**d-f**, Receiver Operating Characteristic (ROC) curves for α, β, and γ cells from the pancreas dataset.

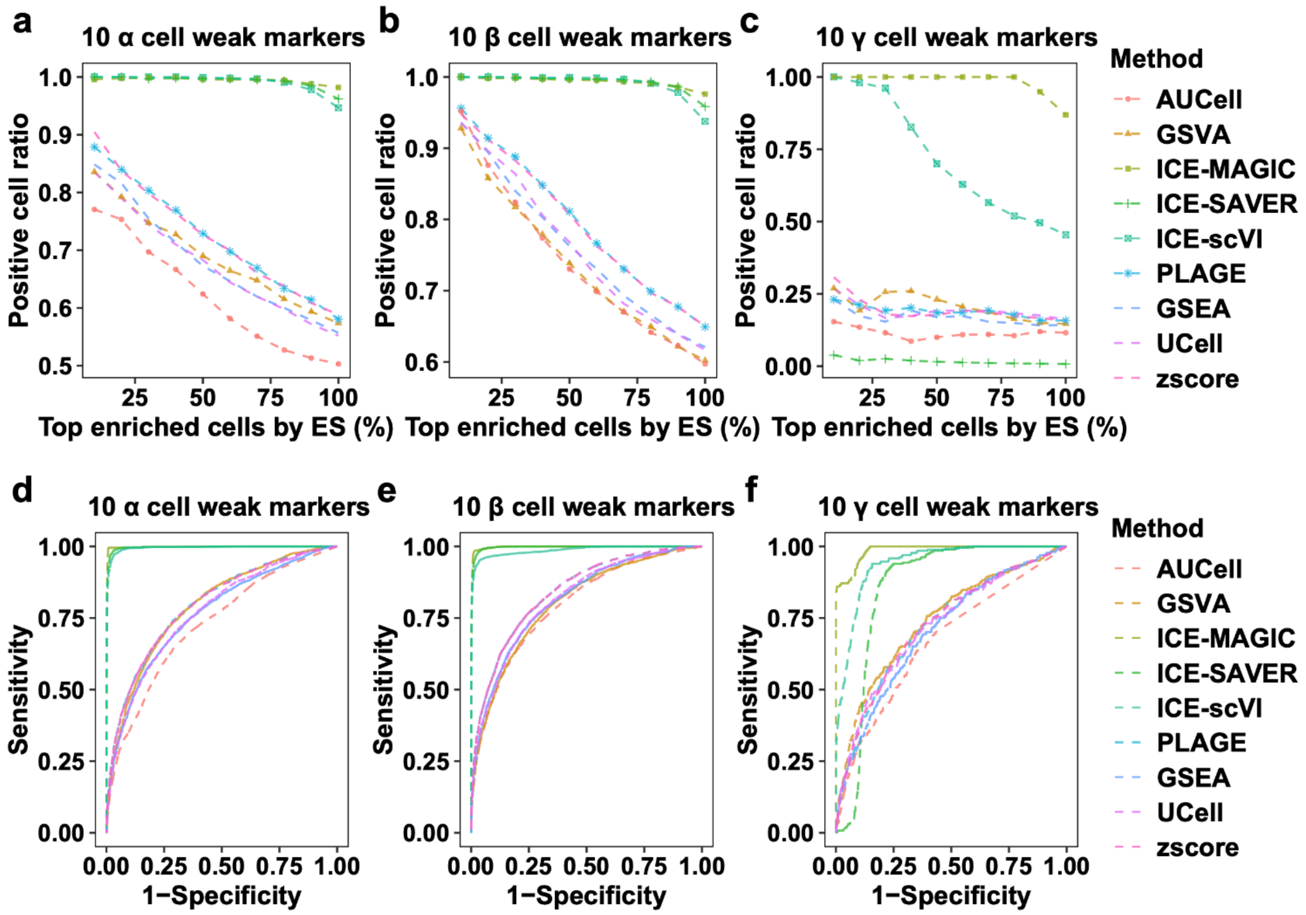


**Fig. S2 | Performance of ICE with different imputation methods compared to other scoring approaches.**

**a-c,** Precision plots across ES-ranked cell intervals using weak markers for α, β, and γ cells. All three ICE variants (MAGIC, SAVER, scVI) achieve higher sensitivity than alternative methods.

**d-f**, Receiver Operating Characteristic (ROC) curves showing the true positive rate versus false positive rate. ICE with MAGIC achieves the best overall performance, though all ICE variants outperform the alternatives.

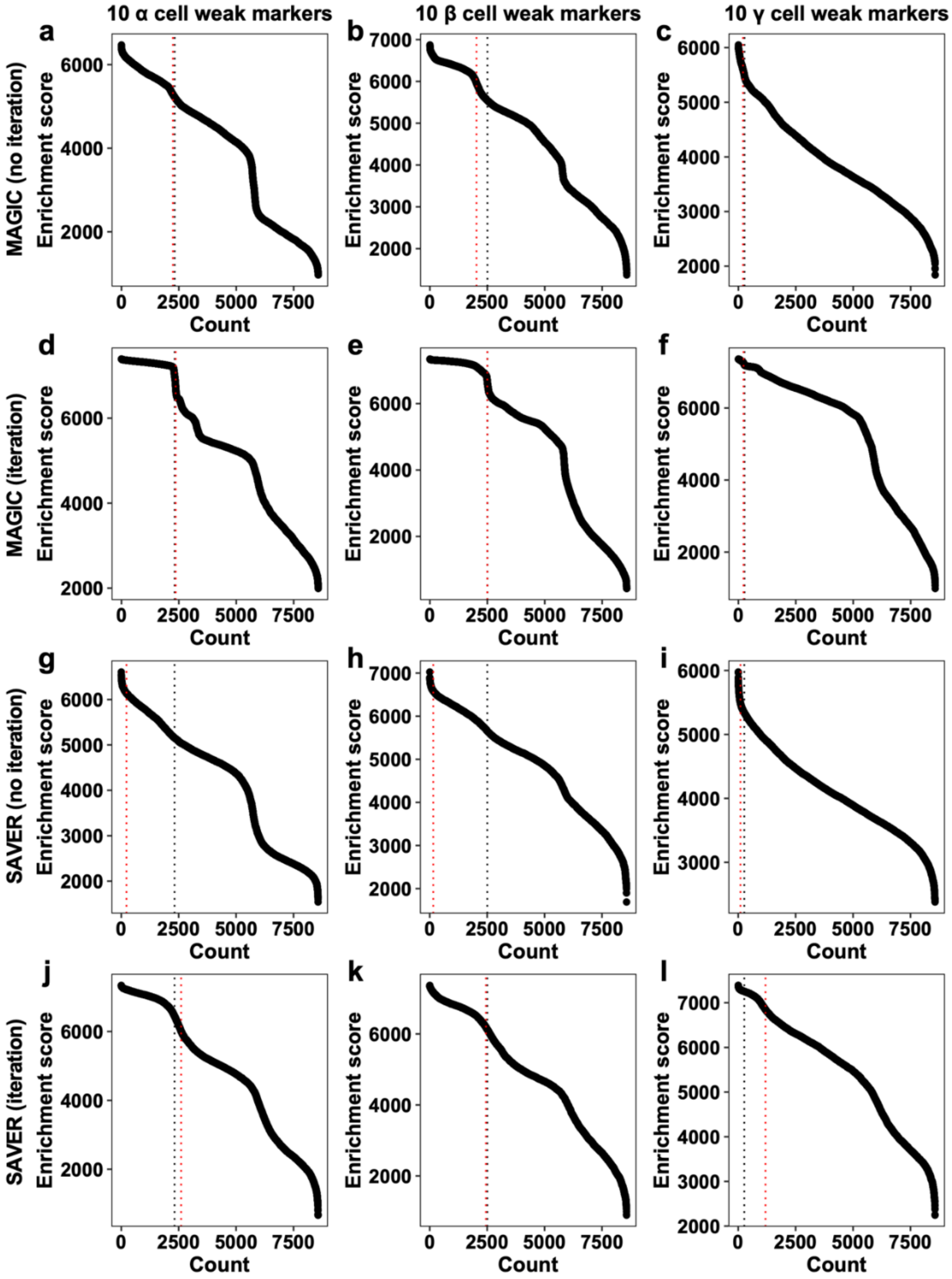


**Fig. S3 | Ablation study demonstrating the distinct contributions of imputation and marker refinement in ICE.** Enrichment Score (ES) distributions for α, β, and γ cells from a pancreas dataset, sorted from highest to lowest. The black dotted line indicates the true cutoff separating cell types, and the red line indicates the cutoff determined by ICE.

**a-c,** MAGIC imputation without marker refinement ("MAGIC no iteration"). Imputation alone correctly identifies α and γ cells but fails for β cells.

**d-f,** Full ICE pipeline with MAGIC ("MAGIC iteration"). Marker refinement enables accurate identification of all three cell types.

**g-i,** SAVER imputation without marker refinement ("SAVER no iteration"), which fails to correctly identify α and β cells.

**j-l,** Full ICE pipeline with SAVER ("SAVER iteration"), where marker refinement improves separation accuracy for α and β cells.

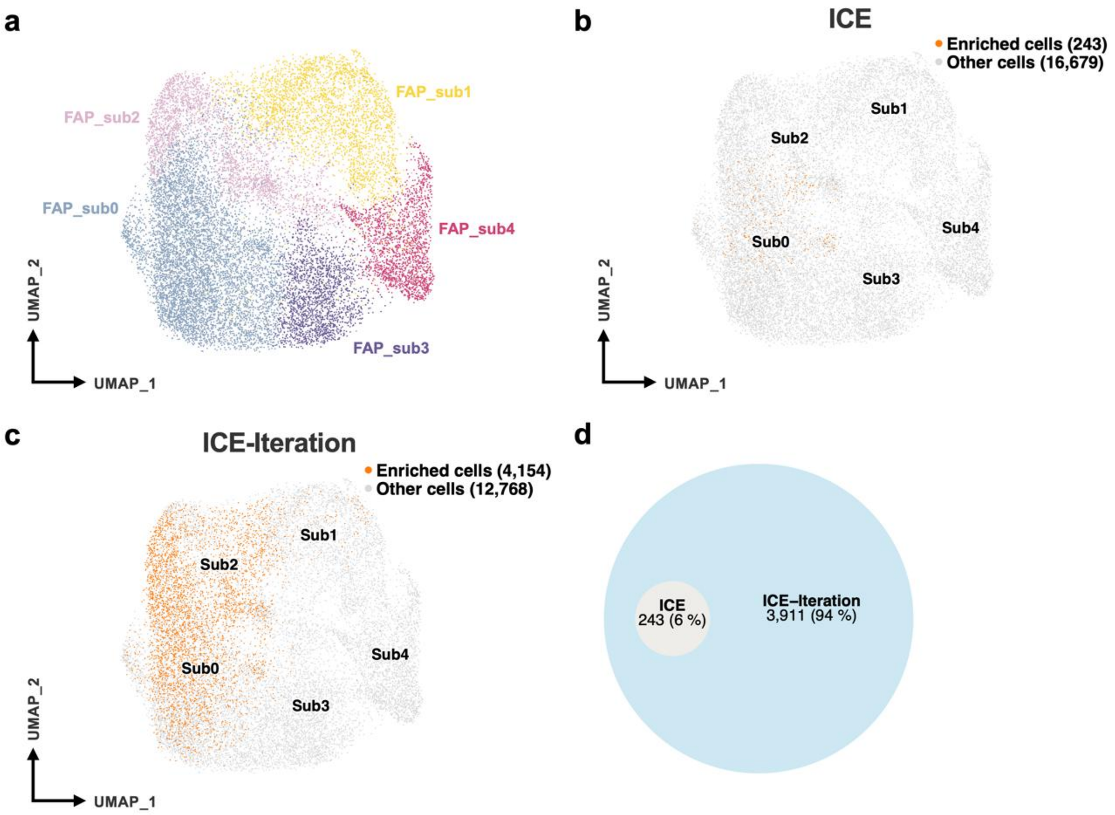


**Fig. S4 | Benchmark analysis of ICE for senescence detection in fibro-adipogenic progenitors (FAPs).**

**a**, UMAP visualization of major clusters in the FAP dataset.

**b**, UMAP showing senescent cells (orange) identified with the SenMayo marker set after imputation but before refinement.

**c**, UMAP showing senescent cells (orange) identified by the complete ICE pipeline with iterative marker refinement ("ICE-iteration").

**d**, Venn diagram comparing the numbers of senescent cells identified by each step. The imputation step identified 243 cells, whereas iterative refinement expanded this population to 4,154 cells, demonstrating the complementary roles of both steps.
